# Temporal shifts in intraspecific and interspecific diet variation: effects of predator body size and identity across seasons in a stream community

**DOI:** 10.1101/476374

**Authors:** Landon P. Falke, Jeremy S. Henderson, Mark Novak, Daniel L. Preston

## Abstract

Intraspecific variation is increasingly recognized as an important factor in ecological interactions, sometimes exceeding the role of interspecific variation. Few studies, however, have examined how intra-versus interspecific variation affect trophic interactions over time within a seasonally dynamic food web. We collected stomach contents from 2028 reticulate sculpin (*Cottus perplexus*), 479 cutthroat trout (*Oncorhynchus clarkii clarkii*), and 107 Pacific giant salamanders (*Dicamptodon tenebrosus*) in western Oregon streams and compared diets among predator species and size classes over three seasons. Predator body size and species identity both showed strong effects on dietary niche breadth, proportional prey composition, and prey size, with seasonal variation in the relative magnitudes of intraspecific and interspecific diet variation. Size-associated diet variation was high in summer and fall but was heavily outweighed by species-associated diet variation in spring. This pattern was driven primarily by a 50-fold increase in the consumption of terrestrial thrips (Order: Thysanoptera) by cutthroat trout in spring compared to summer and fall. Mean dietary niche breadth generally increased with body size and was roughly half as wide in sculpin as in trout and was intermediate in salamanders. Predator-specific responses to the seasonality of terrestrial prey availability were associated with interspecific differences in foraging mode (e.g., benthic vs drift-feeding) and contributed to temporal variation in the roles of predator size and identity in trophic niche differentiation. Our results thereby demonstrate that intraspecific and interspecific diet variation can exhibit strong seasonality in stream predators, emphasizing the dynamic nature of food webs and the need to incorporate sampling over relevant temporal scales in efforts to understand species interactions.

## Introduction

Understanding the relative importance of interspecific versus intraspecific niche variation has long been recognized as being key to understanding the structure and dynamics of communities (May and MacArthur 1972, Shoener 1974, Wiens 1977, Lichstein et al. 2007, Violle et al. 2012, Hart et al. 2016). Nevertheless, ecologists have largely focused the study of interacting populations at the species level, often ignoring the role of variation within species (Abrams and Ginzburg 2000, Bolnick et al. 2011, Novak et al. 2016). Individuals within species often differ from one another in many ecologically meaningful ways, including prey preferences (Estes et al. 2003), microhabitat use (Schlosser 1987), vulnerability to predation (Kusano 1981), and competitive ability (Svanback and Bolnick 2007). For instance, phenotypic changes that occur throughout ontogeny (e.g., body size, physiology, behavior) often alter the types and strengths of interactions in which individuals participate (Polis 1984, Bolnick et al. 2003, Bolnick et al. 2011). Because recent empirical studies have demonstrated that intraspecific variation can influence community and ecosystem processes as much as or even more than interspecific variation (Des Roches et al. 2018), a renewed emphasis is being placed on understanding the mechanisms that drive variation within and among species.

A focal point for research on intraspecific variation has been its influence on species coexistence (Lichstein et al. 2007, Miller and Rudolf 2011, Nakazawa 2015, Bassar et al. 2017). According to the competitive exclusion principle, species cannot stably coexist if they occupy the same ecological niche (Gause 1934, Hutchinson 1957), with only the differential use of resources permitting the coexistence of ecologically similar species (Chesson 2000). Such niche differentiation can be achieved through three basic means: specialization on a distinct set of resources (MacArthur and Levins 1967, Chesson 2000), differential utilization of resources in space (May and Hassell 1981), and differential utilization of resources in time (Armstrong and McGehee 1980, Chesson 1985). Several recent studies have demonstrated the importance of intraspecific variation for community dynamics and the maintenance of species coexistence (Hughes et al. 2008, Clark et al. 2010, Jung et al. 2010, Messier et al. 2010, Bolnick et al. 2011, Pruitt and Ferrari 2011, Violle et al. 2012), yet consensus on the underlying mechanisms and their effects remains elusive. While some theory suggests that intraspecific variation should promote coexistence (Clark et al. 2003, Hubbell 2005, Fridley et al. 2007, Lichstein et al. 2007), other theory suggests that, if anything, individual variation is more likely to prevent species coexistence (Taper and Case 1985, Crutsinger et al. 2008, Hart et al. 2016). Understanding the context for these effects and their underlying mechanisms will benefit from studies that quantify interspecific and intraspecific variation concurrently over space and time in natural communities.

While intraspecific variation is increasingly being quantified with respect to resource use in particular (Araújo et al. 2007, Semmens et al 2009, Coblentz et al. 2017), few studies have considered how the relative magnitudes of intraspecific versus interspecific diet variation may change over time. This is important because failure to consider time scales may bias inferences of the strength and consistency of diet variation within and among species, especially in temporally variable environments (Tinker et al. 2012, Novak and Tinker 2015). For instance, in aquatic communities, which often exhibit considerable within-species variation in body size, interactions between predator and prey can vary greatly over time (Closs and Lake 1994, Dodds et al. 2013, Peralta-Maraver et al. 2017, Heng et al. 2018). This temporal variation in trophic interactions can have strong effects on community dynamics, especially in freshwater streams that have recurrent seasonal changes in community structure, hydrological discharge, primary production, nutrient dynamics, and riparian influences (Closs and Lake 1994, Chaplin et al. 1997, Nakano et al. 1999b, Baxter et al. 2005, Power et al. 2008, Li et al. 2016). Because individuals that differ in species identity and body size typically interact differently with their environment, the relative magnitudes of intra-versus interspecific diet variation in stream food webs may be highly variable over time.

In the present study, we assessed seasonal variation in the interspecific (i.e., taxonomic identity) and intraspecific (i.e., body size) feeding interactions of a stream community. We focused on three generalist predators that co-occur in forested streams throughout western Oregon: reticulate sculpin (*Cottus perplexus*), coastal cutthroat trout (*Oncorhynchus clarkii clarkii*), and Pacific giant salamanders (*Dicamptodon tenebrosus*). We found substantial seasonal variation in the magnitudes of intraspecific and interspecific diet variation among these three focal predators. This resulted primarily due to species-specific responses to seasonal changes in the prey community. Our study thereby demonstrates the importance of considering both forms of variation across time when seeking to understand how trophic interactions influence community dynamics.

## Methods

*Study System* – We examined the stomach contents of reticulate sculpin, cutthroat trout, and Pacific giant salamanders at nine sites in Soap, Oak, and Berry Creeks located within Oregon State University’s McDonald-Dunn Research Forest northwest of Corvallis, Oregon (44.638 N, 123.292 W). Reticulate sculpin are small benthic fishes that prey primarily on benthic macroinvertebrates (Bond 1963, Petrosky and Waters 1975, Wydoski and Whitney 1979, Scott and Crossman 1998, Preston et al. 2017). As aquatic larvae, Pacific giant salamanders are also benthic stream predators that consume a range of prey, including benthic invertebrates, terrestrial arthropods, and other stream-dwelling vertebrates (Kelsey 1995, Cudmore and Bury 2014). In contrast to sculpin and salamanders, trout are active swimmers that feed at both the water surface and the benthos on a relatively even mixture of terrestrial and aquatic prey (Chapman and Bjornn 1969, Jenkins et al. 1970, Elliot 1973, Ware 1973). Our study streams were small (~1 to 3 m in width) and flowed through mixed deciduous-coniferous forests into higher order tributaries of the Willamette River. These streams support a diverse community of aquatic macroinvertebrates (>325 species) (Anderson and Hansen 1987).

*Data Collection* - To collect stomach contents, we conducted electro-fishing surveys in summer (June and July 2015), fall (September 2015), and spring (April 2016). We conducted surveys during the day (0900 – 1700) at three reaches within each stream. Each reach measured ~45 m in length and contained a combination of pool and riffle habitats. To capture predators, a crew of four researchers conducted a single pass of electro-fishing using a backpack electroshocker (Smith-Root LR20B), a block net (1.0 x 1.0 m) and two dip nets (0.30 x 0.25 m). Captured predator individuals were anesthetized, measured for total length, lavaged nonlethally to collect stomach contents using a 60-cc syringe with a blunt 18-gauge needle, and released back into the stream following a recovery period in aerated stream water. Large trout and salamanders were lavaged using a small straw (2.5 mm in diameter) attached to a 500-mL bottle of stream water. We did not lavage individuals smaller than ~25 mm. Stomach contents were preserved in 70% ethanol and later identified in the laboratory using a dissecting microscope (8X to 35X magnification) to the lowest possible taxonomic level (mostly family) according to Merritt et al. (2008). Total lengths of whole, intact prey items were measured to the nearest 0.5 mm. Additional details about study sites and data collection may be obtained in Preston et al. (2018a) and Preston et al. (2018b).

*Data Analyses* – The overall goals of our analyses were to describe and quantify how diets differed by species, predator size, and season. We were especially interested in whether the relative effects of species and size were consistent over time, or whether they varied seasonally. To examine diet variation by predator size, we subdivided each predator species into size classes based on the distribution of total lengths observed when the individuals of all seasons were combined. Sculpin and trout were subdivided into small, medium, and large size-classes using the 25th and 75th percentile of their distributions to ensure similar numbers of individuals in each size class. Due to a relatively low sample size of salamanders, we used the 50th percentile to divide salamanders into two size-classes: small and large.

Among species, size-classes, and seasons, we compared the proportional diet composition by prey counts of seven primary groups: Diptera larvae (true flies), Ephemeroptera larvae (mayflies), Trichoptera larvae (caddisflies), aquatic snails (*Juga sp.*), emergent adult insects (i.e., aquatic insects that have emerged from the stream), terrestrial prey (i.e., organisms with no aquatic life stage), and “other”. The “other” category represented <5% of total prey items. We calculated proportional diet composition in each season by dividing the total number of a given prey group found in the stomach contents of a given species (or size class) by the total number of prey items found in the stomach contents of that species (or size class). Permutational multivariate analysis of variance (PERMANOVA) was performed to assess the statistical strength of differences in individual-level dietary composition due to predator species, body size, and season using the ‘*adonis*’ function in the ‘*vegan*’ R-package (Oksanen 2015). Prey counts were fourth-root transformed prior to PERMANOVA to decrease the effects of extremely large counts of diet items.

We quantified dietary niche breadth at both the species and size-class levels using Levins’ standardized measure (Levins 1968, Hurlbert 1978),

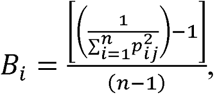

where *p_ij_* is the proportion of prey type *i* in the stomachs of predator group *j* and *n* is the total number of prey items consumed by predator group *j*. Standardized niche breadth values range from 0 (highly specialized) to 1 (highly generalized). Prey items were not grouped into categories for these niche breadth calculations. Instead, we used the lowest possible taxonomic level of prey identification (as suggested by Greene and Jaksic 1983) and treated different prey life stages (i.e., larval, pupal, adult) as distinct prey types. Niche breadth means and standard errors were estimated by non-parametric bootstrapping (1000 draws of 20 prey items) to account for differences in sample sizes among seasons and predator groups. To evaluate diet breadth in the context of prey size, we performed quantile regression analyses (using the 90^th^ and 10^th^ quantiles) on the total lengths of predator individuals and their diet items using the ‘*quantreg*’ package in R (Koenker 2015).

To classify feeding relationships by predator group, we applied a hierarchical cluster analysis on dietary compositions using prey-type proportions and generated dendrograms depicting dietary dissimilarity among size-classes of each predator species within each season. Cluster analyses were performed using the unweighted pair-group method (UPGMA) with Euclidean distance to describe the dissimilarity in dietary composition (Ward 1963, Krebs 1989, Amundsen et al. 2003). To supplement the cluster analyses, we calculated dietary overlap among species and size classes within each season using Schoener’s index of percent overlap (Shoener 1970),

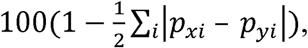

where *p_xi_* and *p_yi_* represent the proportion of prey type *i* in the stomachs of predator species (or size classes) *x* and *y*, respectively. Percent overlap ranges from 0% (no overlap) to 100% (complete overlap). Schoener’s index is considered to be an adequate measure of dietary overlap in the absence of prey availability data (Hurlbert 1978, Wallace 1981).

## Results

We collected stomach contents from a total of 2028 sculpin, 479 trout, and 107 salamanders and found 22,798 identifiable prey items belonging to 104 prey types (entailing taxa and life stages within them). Predator body sizes ranged from 27 to 242 mm in total length, with salamanders and trout reaching considerably larger sizes than sculpin (Fig. 1). Diptera larvae and Ephemeroptera larvae were the primary prey groups found in stomach contents, constituting 36.9% and 36.4% of all food items across all predators, respectively (Fig. 2). Cannibalism was observed in 14 sculpin (13 singletons, 1 doubleton) and 2 trout (both singletons), and predation on sculpin was observed in 2 salamanders (both singletons). We could not find identifiable prey in 115 sculpin (5.67%), 19 trout (3.97%), and 13 salamanders (12.1%) (see Table S1 for numbers of stomachs sampled and percent empty by size class and season).

**Figure 1.**
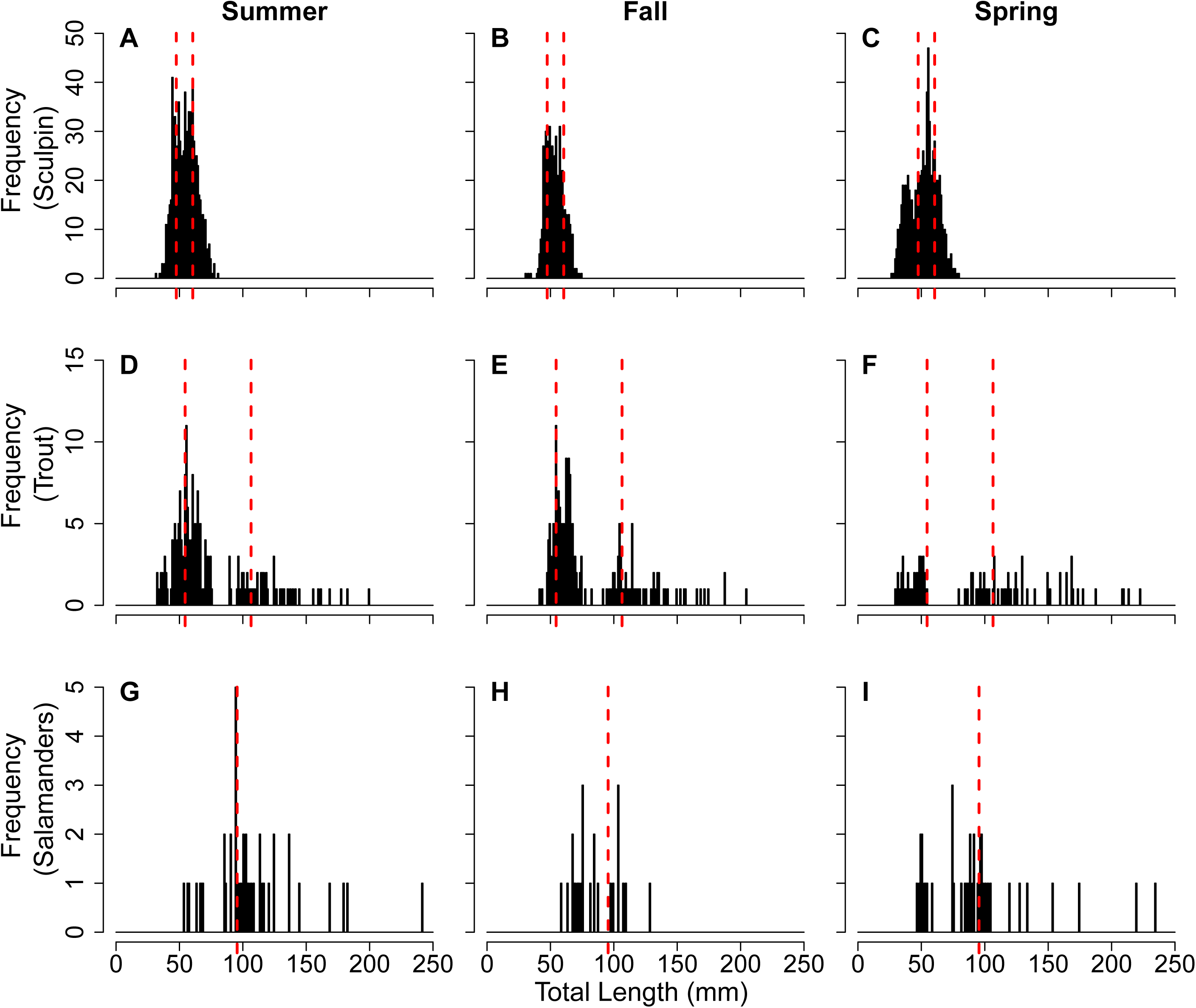
Frequency histograms of total lengths for sculpin (A-C), trout (D-F), and salamanders (G-I) in summer, fall, and spring. Dashed vertical red lines depict the division of size classes (25th and 75th percentile of cumulative size distribution for sculpin and trout; 50th percentile for salamanders).

**Figure 2.**
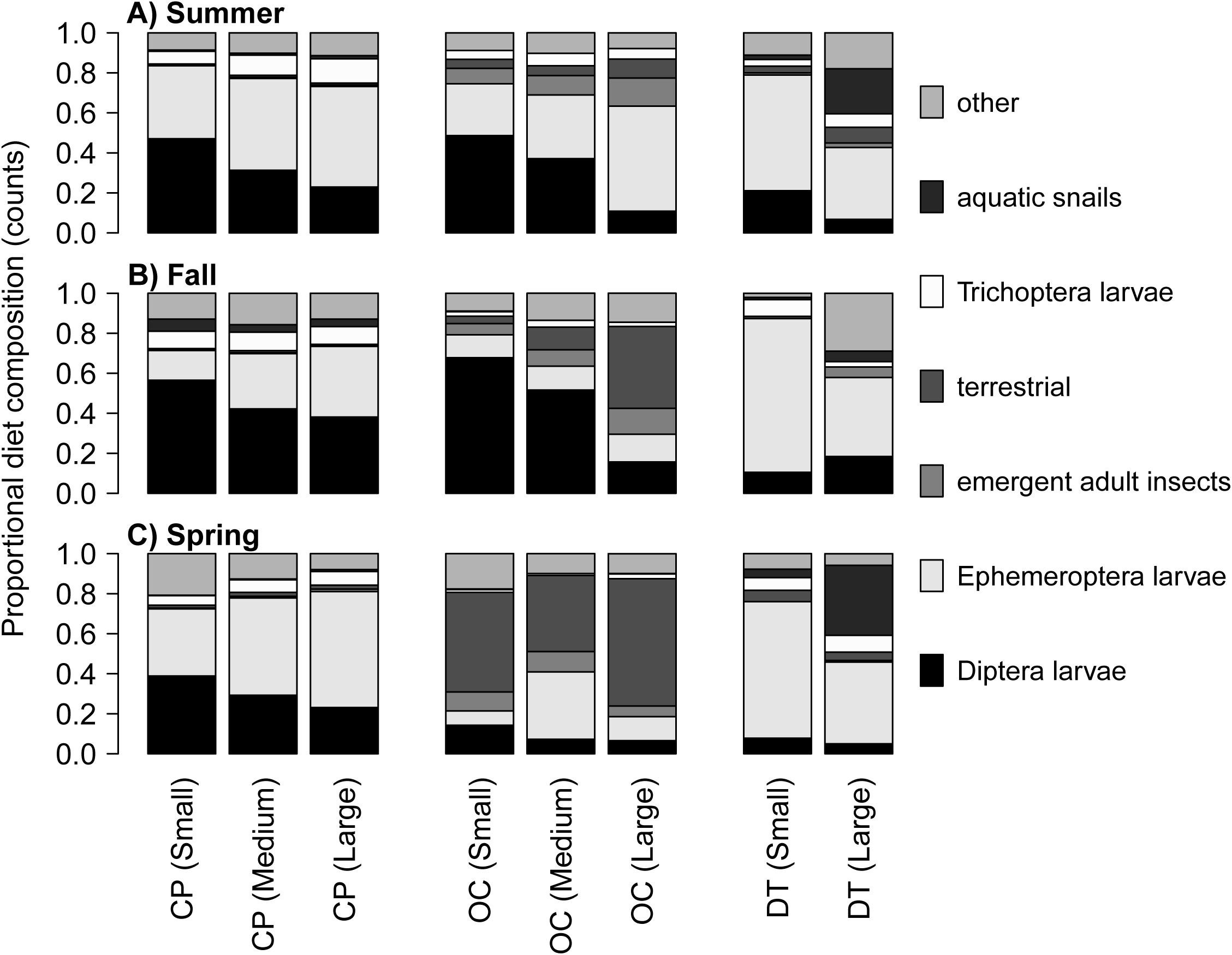
Proportional composition (based on counts) of primary prey groups in diets of sculpin (CP), trout (OC), and salamanders (DT) in summer (A), fall (B), and spring (C). Adult stages of aquatic insects are grouped separately from terrestrial organisms that undergo no aquatic life stages. The “other” category is comprised of aquatic and semi-aquatic prey that individually amounted to less than 5% of predator diets.

In general, sculpin and salamander diets were comprised primarily of benthic aquatic invertebrates, whereas trout diets were comprised of a more even mixture of terrestrial, aquatic, and semi-aquatic prey. Trout diets, which contained the highest overall proportions of adult aquatic insects (9.4%) and terrestrial prey (19.8%), also exhibited the greatest seasonal variation, including a shift in proportional consumption of terrestrial thrips (Order: Thysanoptera) from less than 1% in summer and fall to 49.8% in spring. Proportions of Diptera larvae and Ephemeroptera larvae found in trout stomachs were also highly variable across seasons; trout diets contained relatively high and even proportions of Diptera and Ephemeroptera in summer (~35% and ~33%, respectively), high proportions of Diptera (~52%) and low proportions of Ephemeroptera (12%) in fall, and low proportions of both in spring (~9% and ~15%) (Fig. 2). Sculpin and salamander diets exhibited relatively minimal seasonal variation in proportional diet compositions compare to trout (Tables S2-S4).

Dietary niche breadth varied seasonally within species but was highest (most generalized) in trout, lowest (most specialized) in sculpin, and consistently higher in the larger size-classes within each species (Fig. 3). Mean dietary niche breadth was roughly half as high in sculpin (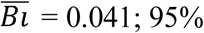 CI = 0.023 to 0.059) as in trout (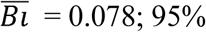 CI = 0.043 to 0.113) over all seasons combined, with salamanders exhibiting intermediate values (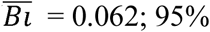 CI = 0.033 to 0.091). Sculpin dietary niche breadth was lowest in summer (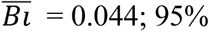 CI = 0.019 to 0.049) and highest in spring (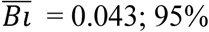 CI = 0.028 to 0.058). Salamander dietary niche breadth was also lowest in summer (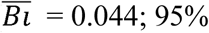 CI = 0.029 to 0.059) but was highest in fall (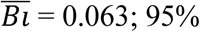 CI = 0.036 to 0.090). Trout dietary niche breadth was lowest in fall (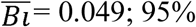 CI = 0.023 to 0.075) and highest in spring (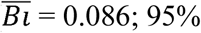 CI = 0.049 to 0.123).

**Figure 3.**
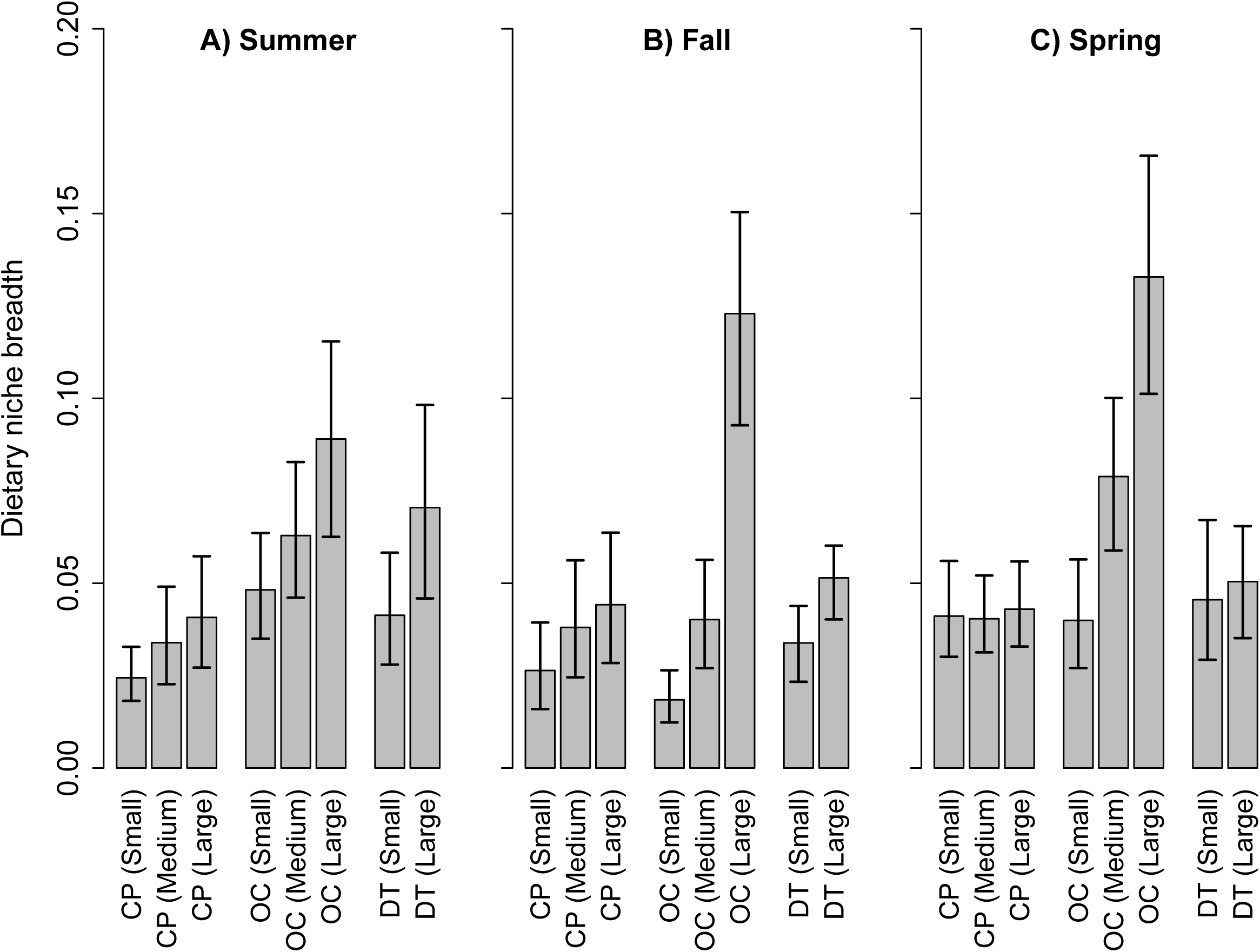
Mean dietary niche breadth (Levins’ standardized measure) by size class for sculpin (CP), trout (OC), and salamanders (DT) in summer (A), fall (B), and spring (C). Lower niche breadth values indicate more specialized (less diverse) diets. Error bars represent 95% confidence intervals around the mean.

Predators of differing body size and of differing identity both differed in the mean size and composition of prey they consumed (Figs 3-4; Tables S2-S4). Mean prey length 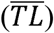 was much larger in salamanders (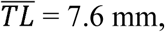, σ = 6.7) compared to sculpin (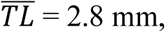, σ = 2.7) and trout (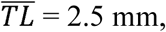, σ = 2.7) and generally increased with increasing predator body size (Fig. 4). For sculpin and trout, Diptera larvae were generally found in higher proportions in the diets of smaller size classes, whereas the proportions of Ephemeroptera larvae were higher in larger size classes. Based on proportional diet composition, large sculpin consumed 19.1% (averaged difference across seasons) fewer Diptera larvae and 19.5% more Ephemeroptera larvae than small sculpin. Similarly, large trout consumed 27.4% fewer Diptera larvae and 11.3% more Ephemeroptera larvae than small trout. Larger trout also consumed higher proportions of adult aquatic insects and terrestrial prey than smaller trout. Small salamander diets contained higher proportions of Ephemeroptera larvae but larger salamanders generally consumed more prey that were not Ephemeroptera of Diptera (e.g., snails, crayfish, annelids, other rare prey). PERMANOVA analysis confirmed the statistical strength of these dietary differences among predator species (pseudo-F = 37.95, p<0.001), body sizes (pseudo-F = 58.23, p<0.001), and season (pseudo-F = 73.57, p<0.001; Table 1).

**Figure 4.**
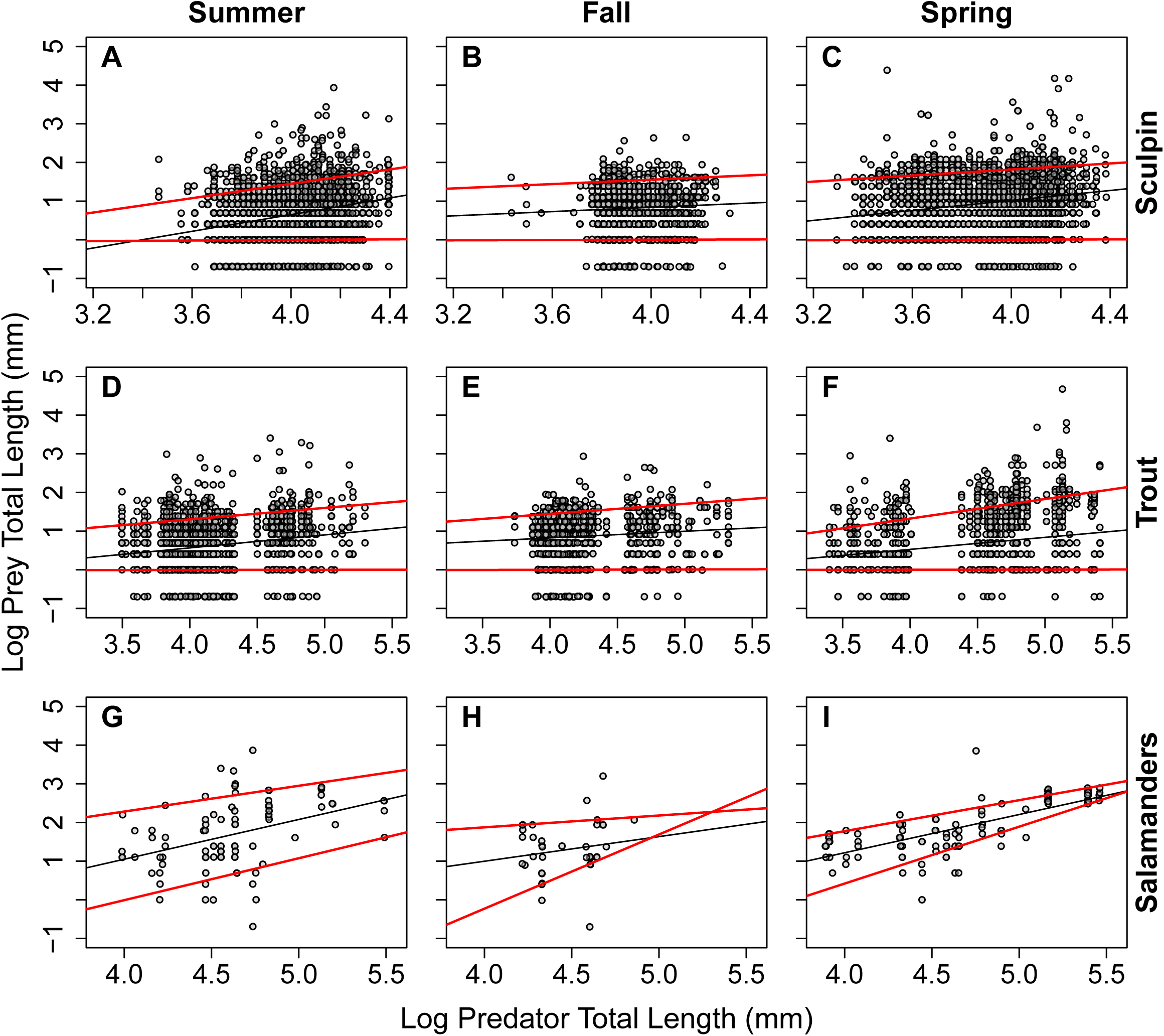
Regression plots depicting the 10th and 90th quantiles for the relationships between total lengths of sculpin (A-C), trout (D-F), and salamanders (G-I) and whole, intact food items in their stomach contents in summer, fall, and spring.

**Table 1.**
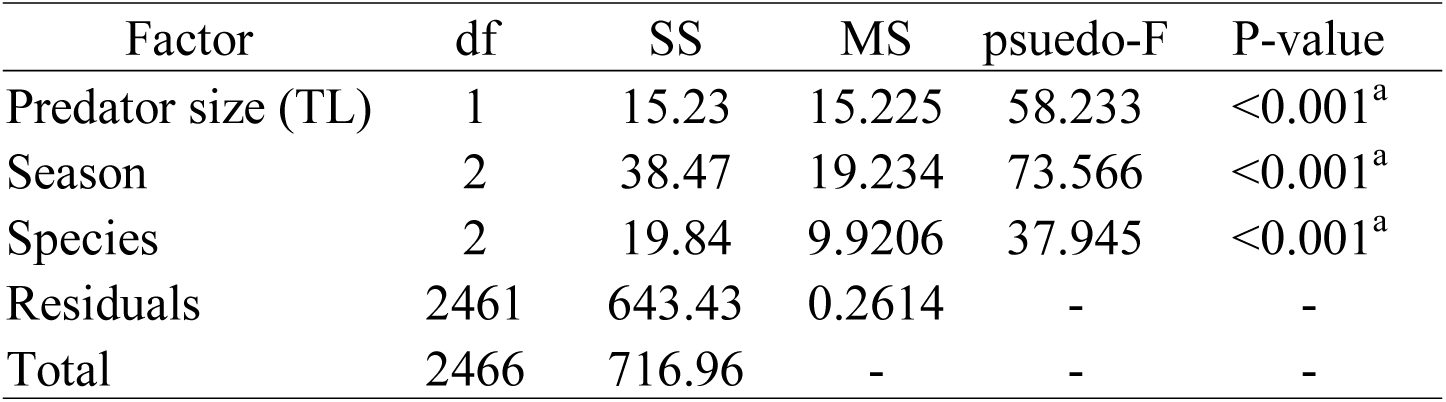
Permutational multivariate analysis of variance (PERMANOVA) of dietary composition with stomachs as sites and predator size, season, and species as fixed factors.

The hierarchical clustering of size classes by diet dissimilarity varied across seasons (Fig. 5). Diet dissimilarity among size classes was lowest in summer (Fig. 5A), intermediate in fall (Fig. 5B), and highest in spring (Fig. 5C). In summer and fall, the diets of heterospecific size-classes were often more similar than the diets of conspecific size-classes. Only in spring were size classes clustered according to predator species, with sculpin and salamander diets being highly dissimilar to trout diets, a result that was driven primarily by a large increase in terrestrial thrips found in trout stomachs. Calculations of percent overlap were consistent with results of the cluster analyses (Table S3). Dietary overlap generally decreased with increased differences in body size within and between species, and trout size-classes showed the lowest dietary overlap with sculpin and salamander size-classes in spring.

**Figure 5.**
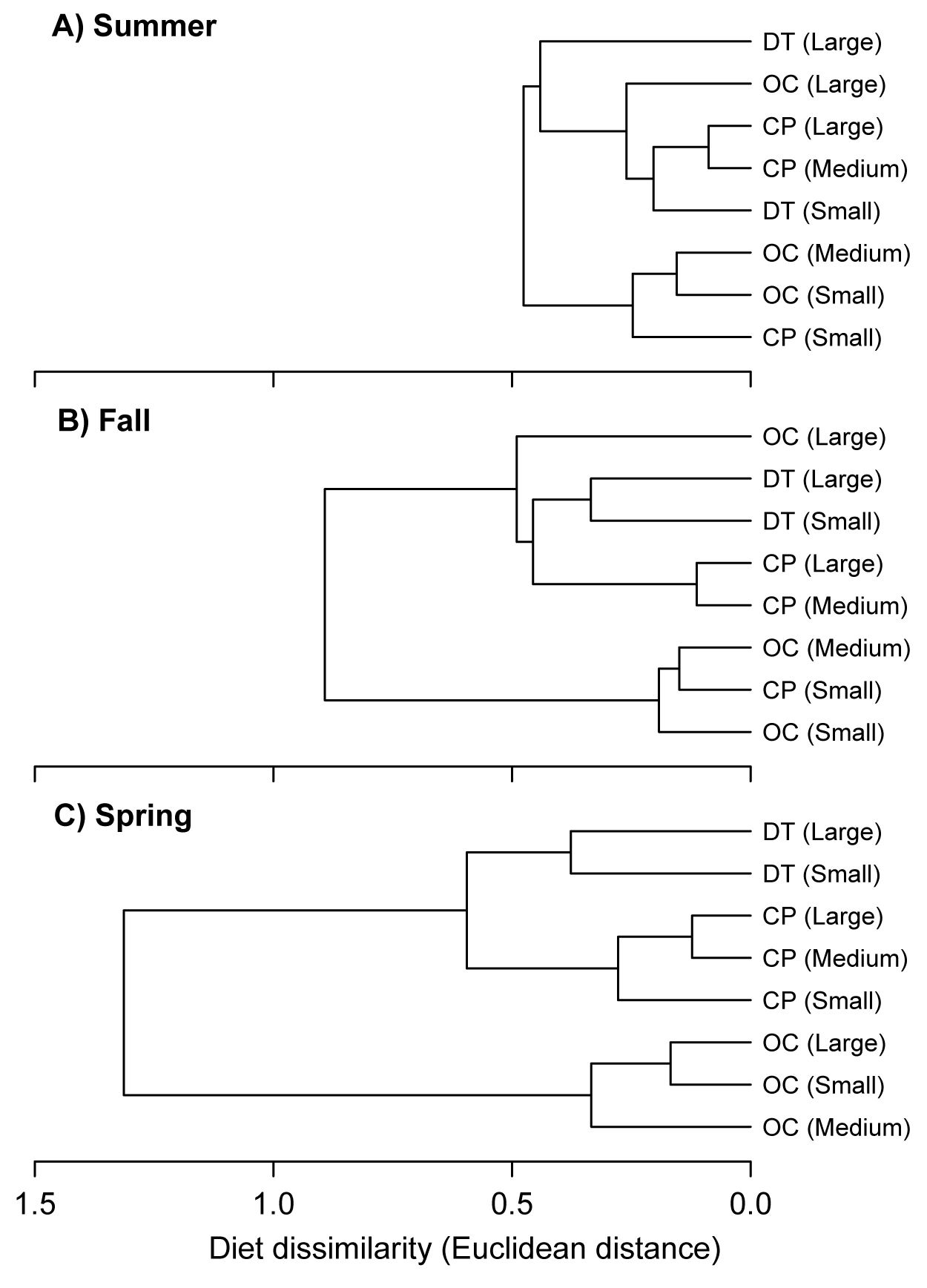
Dendrograms generated from hierarchical cluster analyses of proportional diet composition in size classes of sculpin (CP), trout (OC), and salamanders (DT) in summer (A), fall (B), and spring (C). Shorter branches represent greater similarity in dietary composition between the connected groups.

## Discussion

The relative magnitudes of intra- and interspecific diet variation are recognized as being important to shaping community dynamics, yet few studies have evaluated the degree to which these two types of variation are static or dynamic in time. In the present study, we compared intraspecific and interspecific variation in feeding interactions within a stream community over three seasons. Our primary finding is that size-associated diet variation had a strong influence on feeding relationships in summer and fall but was heavily outweighed by species-associated diet variation in spring. This pattern emerged largely due to predator-specific responses to seasonal changes in prey availability. Our study therefore demonstrates that the relative roles of intraspecific and interspecific variation can be temporally dynamic, thereby emphasizing the importance of considering them at relevant temporal scales, beyond just ‘snapshots’ in time.

Seasonal changes in prey composition and availability contributed to the temporal variation in intra- and interspecific diet variation. Trout diets were comprised of a mixture of terrestrial and aquatic prey but were highly variable across seasons, with spring diets containing high proportions of terrestrial prey. In contrast, sculpin and salamander diets were comprised primarily of benthic aquatic prey and showed relatively low seasonal variation. While seasonal inputs of terrestrial invertebrates can have strong effects on trophic interactions in streams (Nakano et al. 1999b, Nakano and Murakami 2001, Kawaguchi et al. 2003, Baxter et al. 2005), their relative effects can differ strongly across predator-prey interactions. For instance, seasonal diet shifts are ubiquitous among freshwater salmonids but are not widely observed in sculpin or salamanders (Wilhelm et al. 1999, Li et al. 2016, Cochran-Biederman and Vondracek 2017). Preston et al. (2018a) also suggest that prey-specific sculpin feeding rates are relatively consistent across space and time in our study streams, especially compared to the more variable trout diets observed in the present study.

Differences in dietary niche breadth and in responses to changing prey availability among our focal predators likely stem from differences in foraging strategies. Sculpin and salamanders are bottom-dwellers that employ ambush predation to feed primarily on benthic macroinvertebrates (Bond 1963, Daniels and Moyle 1978, Kratz and Vinyard 1981, Wells 2007, Cudmore and Bury 2014). In contrast, trout are active swimmers that exhibit a wider diet breadth because they feed on both aquatic and terrestrial prey in the benthos and throughout the water column (Chapman and Bjornn 1969, Jenkins et al. 1970, Elliot 1973). Trout diets are therefore expected to exhibit greater responses to the changes in availability of terrestrial prey, that are inherent to streams, than are the diets of sculpin and salamanders. Because of the high seasonal variation in availability of terrestrial prey compared to the relative consistency of the benthic prey community in our study streams (Preston et al. 2018a), trout diets showed much greater seasonal variation compared to sculpin and salamander diets. These results are consistent with prior experimental work. By manipulating inputs of terrestrial invertebrates into streams, Gillette (2012) demonstrated that stream predators can indeed exhibit species-specific responses to changes in the availability of terrestrial prey based on differences in diet breadth and foraging behavior. Species-specific responses are also observed in riparian consumers when aquatic derived subsidies are manipulated (Paetzold et al. 2006, Marczak and Richardson 2007). Taken together, these results emphasize how the interplay of predator characteristics (e.g., diet breadth and foraging behavior) and prey characteristics (e.g., seasonal changes in availability) can interact to drive temporal shifts in interspecific diet variation.

The high seasonal diet variation of trout was driven largely by consumption of western flower thrips (Thysanoptera: Thripidae), which comprised nearly half of all prey consumed by trout in the spring but comprised less than 1.0% in summer and fall. Thrips were not a major diet item in sculpin or salamanders in any season. Thrips, which are considered widespread agricultural pests (Teulon et al. 1993), hatch from eggs in plant tissues and later drop to the ground before metamorphosing into adults (Sanderson 1990). The timing of the thrip lifecycle is temperature dependent, with the larval stage lasting just five to 20 days. Thus, larval thrips apparently dropped from overhanging canopy into our study streams prior to or during the spring surveys and were subsequently consumed at disproportionately high rates by trout compared to the consumption rates by sculpin and salamanders. Previous evidence of thrip consumption by stream predators is scarce but has been documented in brook trout (Williams 1981) and sticklebacks (Hynes 1950). However, similar temporally-pulsed inputs of terrestrial arthropods are well documented in forested streams and are known to contribute a large portion of available prey for streams predators (Wipfli 1997, Kawaguchi and Nakano 2001, Romero et al. 2005, Chan et al. 2007). Consumption of cross-ecosystem subsidies by top predators such as trout can reshape the structure and energetic dynamics of stream food webs (Perkins et al. 2018). Our results exemplify how specific community members (i.e., thrips and cutthroat trout in our study) can disproportionally contribute to cross-ecosystem fluxes of nutrients and matter from land to water (Polis et al. 2004).

Predator-prey body sizes had a strong influence on feeding relationships, especially in summer and fall. For instance, salamanders, which exhibited some of the largest body sizes among our focal predators, fed on substantially larger prey, including prey taxa that were rare in sculpin and trout diets (e.g., crayfish, caterpillars, snails). Despite aquatic snails (*Juga* sp.) being the most abundant benthic macroinvertebrates by biomass in our study streams (Preston et al. 2018b, Hawkins and Furnish 1987), sculpin feeding rates on snails were among the lowest for all observed prey, likely reflecting their low digestibility (Preston et al. 2018b) and morphological constraints of predators (e.g., gape width and size of digestive tract). Additionally, cannibalism and intraguild predation among our focal predators (i.e., sculpin eating sculpin, trout eating trout, salamanders eating sculpin) was more common in larger individuals. The influence of body size on feeding interactions is not surprising given that body size is widely recognized as a key trait dictating an organism’s trophic ecology and interactions with its environment (Werner and Gilliam 1984, Woodward et al. 2005, Woodward and Warren 2005, Petchey et al. 2008, Rudolf et al. 2014). For example, Woodward and Hildrew (2002) demonstrated that feeding relationships within a guild of stream predators were driven primarily by body-size constraints that led to less diverse diets in smaller predators by restricting consumption to a limited subset of the prey-size spectrum. Consistent with this result, we found that both dietary niche breadth and prey size generally increased with predator body size, and dietary overlap was generally higher between groups of similar body size. In several cases, dietary overlap was higher between heterospecifics of similar body size than between conspecifics of dissimilar body size (see Table S5). In summer and fall, for example, higher dietary overlap was observed between small trout and small sculpin than between small sculpin and large sculpin. In spring, however, dietary overlap between small sculpin and small trout was less than half the overlap between small and large sculpin, which again reflects the influence of terrestrial prey availability on feeding relationships in our study.

Temporal variability in the magnitudes of intra-versus interspecific variation may play an important role in the coexistence of our focal predators by limiting similarity in resource use over time. According to classic niche theory, increased dissimilarity in resource use should lead to decreased competition (MacArthur and Levins 1967, File et al. 2012). Thus, seasonal shifts in intra- and interspecific diet variation, such as those caused by changes in prey availability, should theoretically coincide with changes in the strength of competition (Zaret and Rand 1971, Chase and Leibold 2003, Correa and Winemiller 2014, Neves et al. 2018). For instance, Zaret and Rand (1971) suggest that increased interspecific competition during the dry season when food resources in tropical streams are low provides a good explanation for seasonal diet shifts in characin fishes. This flexibility of resource partitioning has been suggested as a key mechanism for coexistence of stream predators (Nakano et al. 1999a, Dineen et al. 2007). Assuming classic niche theory holds true, if interspecific diet variation is lower (i.e., higher species-level overlap in diet) at times of the year when terrestrial prey inputs are scarce, then intraspecific diet variation may provide a seasonally important mechanism to promote species coexistence by reducing species-level resource overlap among our focal predators. However, because (1) we did not estimate feeding rates nor quantify the limiting availability of prey, (2) species overlapping in resources do not necessarily compete (Menge 1979), (3) competition may occur along multiple niche dimensions beyond diet (Pianka 1975), and (4) patterns of niche overlap may reflect “ghosts of competition past” (Connell 1980), we are here unable to infer the strength of competition in our system. Our results nevertheless provide strong evidence that the relative magnitudes of intra- and interspecific diet variation can change over time, and hence competition is likely to exhibit temporal variation as well. Our study therefore suggests that temporal scales are an important consideration in efforts to understand coexistence.

Given that seasonality in environmental factors and the strength of predator-prey interactions is widespread across various ecosystems and taxa (Ostfeld and Keesing 2000, Thompson et al. 2012, Humphries et al. 2017, Calizza et al. 2018), temporal variation in the relative magnitudes of intraspecific and interspecific diet variation is likely to be inherent to most food webs. The seasonal influx of a single prey type, terrestrial thrips, into our study streams was enough to substantially increase interspecific diet variation in spring. Examples of temporal pulses in the availability of even a single prey type are widespread and occur across various time scales (Yang et al. 2010): diurnal pulses of marine copepods consumed by pelagic fishes (Godin 1981), annual pulses of anadromous fish carcasses providing food for minks (Ben-David 1997), and multiannual fluctuations in abundances of rodents consumed by owls (Korpim□ki 1992). Diet studies conducted on time scales that are poorly matched to the relevant intrinsic and extrinsic drivers, such as seasonal variation in prey communities, may not capture the full picture of how temporally dynamic trophic interactions can be in nature. In the present study, our estimates of diet variation are averaged over multiple weeks and compared across seasons, whereas higher (or lower) diet variation may be revealed on much different time scales (e.g., diurnal, annual, decadal). We recommend that future studies of trophic interactions should consider temporal changes in prey populations and incorporate time scales that are relevant to the life histories of the interacting species.

## Supporting information

## Acknowledgements

For assistance with data collection we thank Anuwar Azraf, Madeleine Barrett, Alicen Billings, Andrew Branka, Clarissa Chan, Rebecca Crawford, Kila Gebeyessa, Daniel Gradison, Kurt Ingeman, Emily Hiser, Tamara Layden, Dana Moore, Elora Ormand, Arren Padgett, Zachary Randell, Kieryian Rock, Wendy Saepharn, Alex Scharfstein, Leah Sequi, Johnny Schwartz, Isaac Shepard, Jasper Shults, Samantha Sturman, Ernesto Vaca Jr, and Beatriz Werber. Input on identifying invertebrates was provided by Michael Bogan, Richard Van Driesche, William Gerth, Stanley Gregory, Judy Li, David Maddison, and Christopher Marshall. Additional feedback on the manuscript was provided by Kyle Coblentz, Shumpei Maruyama, and Zachary Randell. We thank the Oregon State University College of Forestry for providing access to field sites in the McDonald-Dunn Research Forest. Funding was provided by the National Science Foundation Grant DEB-1353827 and Oregon State University.

