## Supplementary material for "Temporal shifts in intraspecific and interspecific diet variation: effects of predator body size and identity across seasons in a stream community"

**Table S1.** Number of stomachs sampled (N) and % empty by season and size class for each predator.

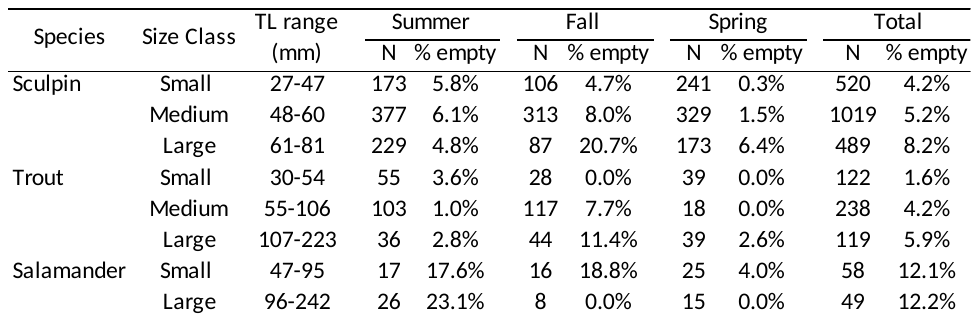

**Table S2.** Summary of proportional diet composition for size classes of each predator in summer.

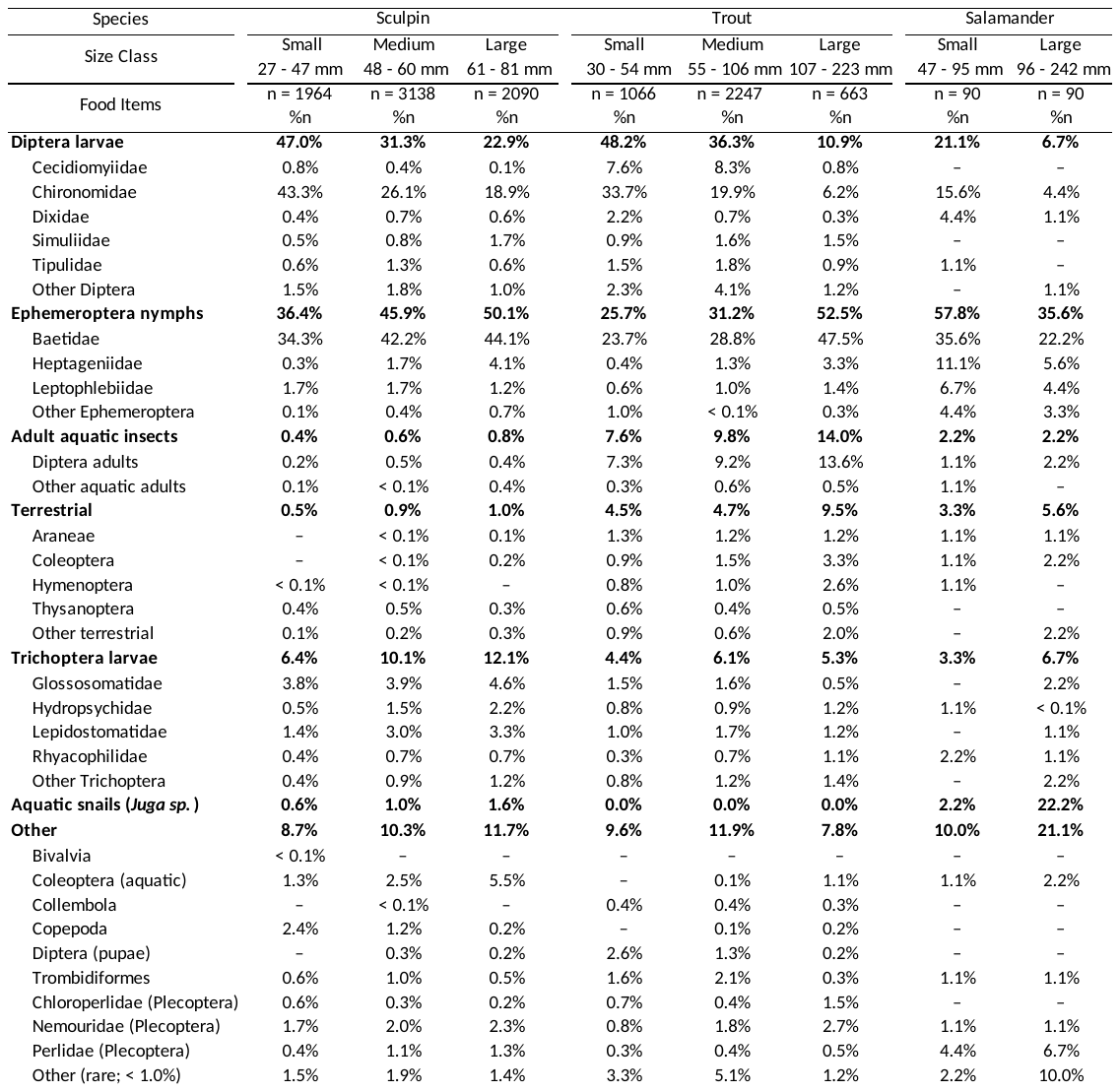

**Table S3.** Summary of proportional diet composition for size classes of each predator in fall.

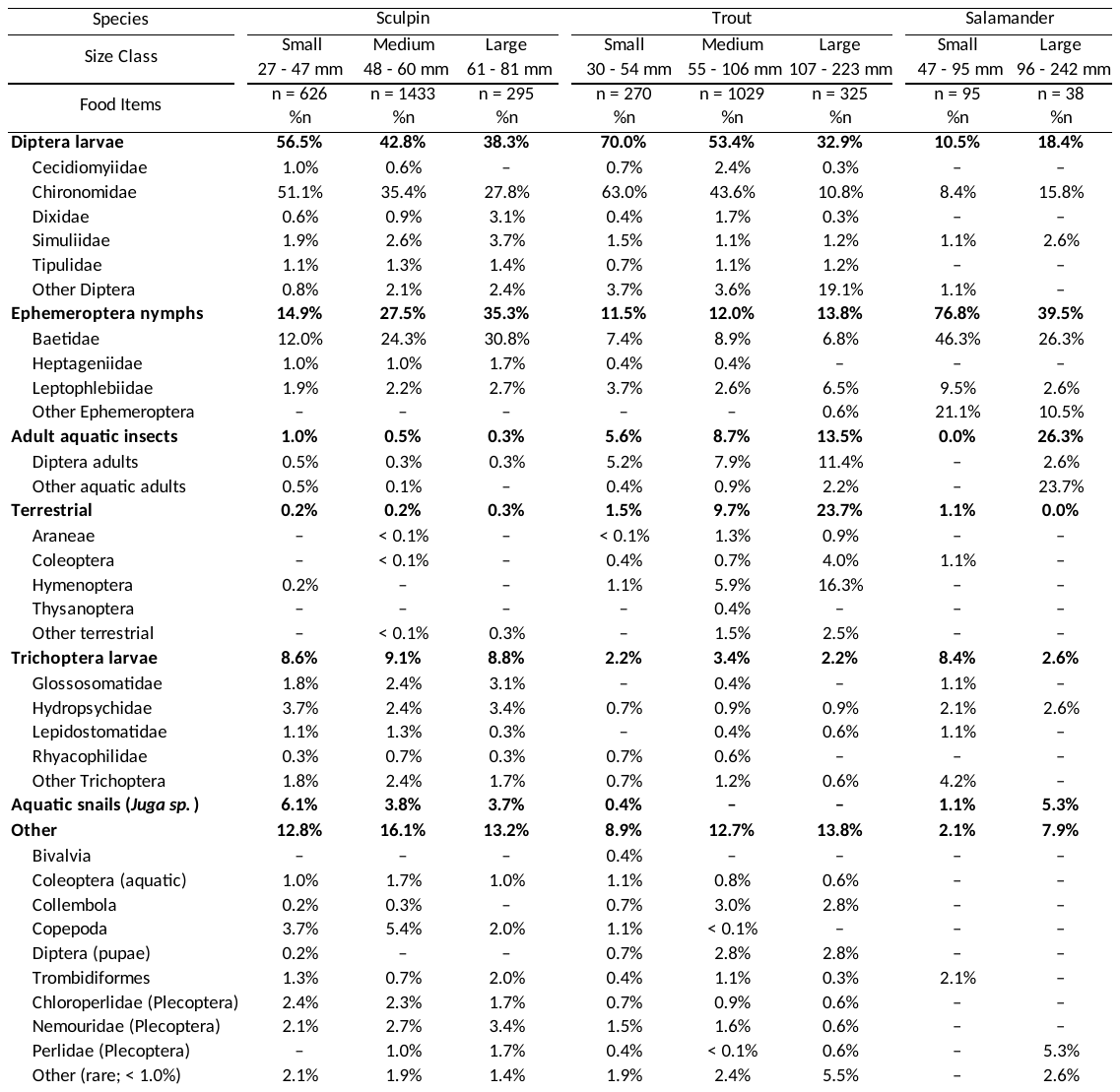

**Table S4.** Summary of proportional diet composition for size classes of each predator in spring.

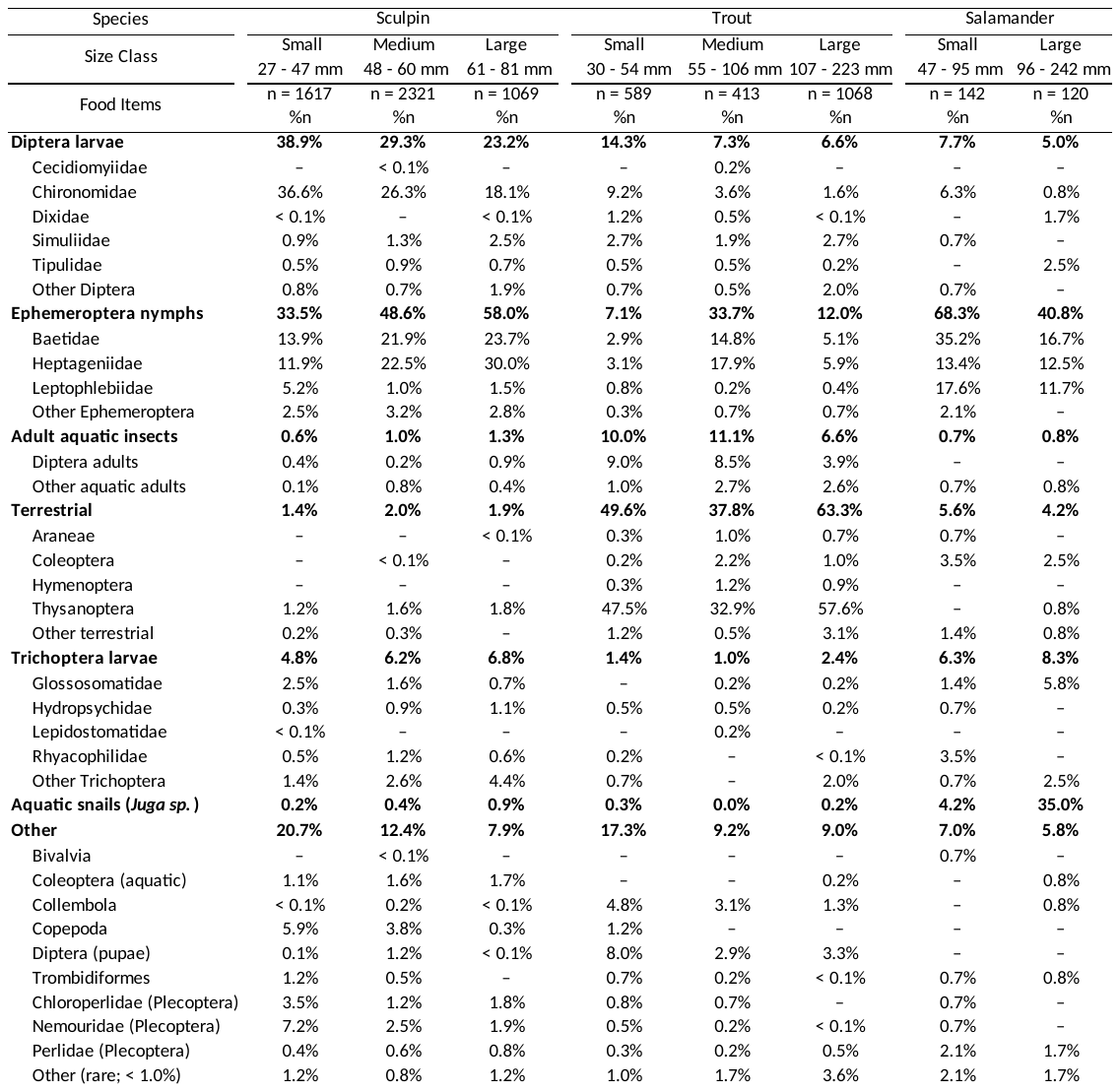

**Table S5.** Percent overlap in diet between size classes of sculpin (CP), trout (OC), and salamanders (DT) in each season.

| Size class combination | | |  | Percent overlap in diet | | | | |
| --- | --- | --- | --- | --- | --- | --- | --- | --- |
|  |  |  |  | Summer |  | Fall |  | Spring |
| CP (Small) | vs | CP (Medium) |  | 72.8% |  | 68.4% |  | 67.4% |
| CP (Small) | vs | CP (Large) |  | 64.6% |  | 60.2% |  | 56.4% |
| CP (Small) | vs | OC (Small) |  | 65.4% |  | 69.6% |  | 24.0% |
| CP (Small) | vs | OC (Medium) |  | 57.8% |  | 63.1% |  | 33.3% |
| CP (Small) | vs | OC (Large) |  | 43.4% |  | 26.3% |  | 22.2% |
| CP (Small) | vs | DT (Small) |  | 54.3% |  | 29.6% |  | 41.5% |
| CP (Small) | vs | DT (Large) |  | 34.7% |  | 38.7% |  | 35.3% |
| CP (Medium) | vs | CP (Large) |  | 73.8% |  | 72.5% |  | 74.5% |
| CP (Medium) | vs | OC (Small) |  | 61.6% |  | 55.1% |  | 24.0% |
| CP (Medium) | vs | OC (Medium) |  | 62.3% |  | 56.1% |  | 42.6% |
| CP (Medium) | vs | OC (Large) |  | 52.0% |  | 27.6% |  | 26.0% |
| CP (Medium) | vs | DT (Small) |  | 61.0% |  | 42.4% |  | 49.1% |
| CP (Medium) | vs | DT (Large) |  | 40.5% |  | 48.9% |  | 37.1% |
| CP (Large) | vs | OC (Small) |  | 54.5% |  | 46.6% |  | 24.8% |
| CP (Large) | vs | OC (Medium) |  | 59.4% |  | 49.2% |  | 43.3% |
| CP (Large) | vs | OC (Large) |  | 53.0% |  | 27.2% |  | 27.6% |
| CP (Large) | vs | DT (Small) |  | 62.1% |  | 49.5% |  | 50.9% |
| CP (Large) | vs | DT (Large) |  | 43.6% |  | 52.1% |  | 38.2% |
| OC (Small) | vs | OC (Medium) |  | 72.1% |  | 68.5% |  | 59.4% |
| OC (Small) | vs | OC (Large) |  | 46.5% |  | 36.0% |  | 53.6% |
| OC (Small) | vs | DT (Small) |  | 50.5% |  | 22.5% |  | 17.4% |
| OC (Small) | vs | DT (Large) |  | 37.2% |  | 31.8% |  | 13.4% |
| OC (Medium) | vs | OC (Large) |  | 55.5% |  | 47.8% |  | 53.5% |
| OC (Medium) | vs | DT (Small) |  | 54.6% |  | 24.5% |  | 36.9% |
| OC (Medium) | vs | DT (Large) |  | 39.6% |  | 32.5% |  | 31.7% |
| OC (Large) | vs | DT (Small) |  | 47.7% |  | 24.1% |  | 21.9% |
| OC (Large) | vs | DT (Large) |  | 38.8% |  | 26.9% |  | 22.3% |
| DT (Small) | vs | DT (Large) |  | 51.2% |  | 51.5% |  | 49.8% |
